# FoxM1 drives proximal tubule proliferation during repair from acute kidney injury

**DOI:** 10.1101/436576

**Authors:** Monica Chang-Panesso, Farid F. Kadyrov, Matthew Lalli, Haojia Wu, Shiyo Ikeda, Akio Kobayashi, Benjamin D. Humphreys

## Abstract

The proximal tubule has a remarkable capacity for repair after acute injury but the cellular lineage and molecular mechanisms underlying this repair response have been poorly characterized. Here, we developed a Kim-1-GFPCreER^t2^ knockin mouse line (Kim-1-GCE), performed genetic lineage analysis after injury and measured the cellular transcriptome of proximal tubule during repair. Acutely injured genetically labeled clones co-expressed Kim-1, Vimentin, Sox9 and Ki67, indicating a dedifferentiated and proliferative state. Clonal analysis revealed clonal expansion of Kim-1+ cells, indicating that acutely injured, dedifferentiated proximal tubule cells account for repair rather than a fixed tubular progenitor. Translational profiling during injury and repair revealed signatures of both successful and unsuccessful maladaptive repair. The transcription factor FoxM1 was induced early in injury, was required for epithelial proliferation, and was dependent on epidermal growth factor receptor (EGFR) stimulation. In conclusion, dedifferentiated proximal tubule cells effect proximal tubule repair and we reveal a novel EGFR-FoxM1-dependent signaling pathway that drives proliferative repair after injury.

## Introduction

Acute kidney injury (AKI) has a wide spectrum of outcomes ranging from full recovery to failed repair and transition to chronic kidney disease. According to a recent report from the CDC examining trends in hospitalizations for acute kidney injury in the US from 2000 to 2014, the rate of AKI hospitalizations increased by 230% over this time frame, going from 3.5 to 11.7 per 1000 persons (1). Furthermore, it has been reported that Medicare patients aged 66 years and older who were hospitalized for AKI had a 35% cumulative probability of a recurrent AKI hospitalization within one year and 28% were diagnosed as having CKD in the year following an AKI hospitalization (2). These troubling statistics points toward a pressing need to identify therapeutic interventions to prevent and treat AKI.

The proximal tubular epithelium makes up the bulk of the kidney cortex and is responsible for reabsorption of a large portion of the glomerular filtered load in order to maintain solute and volume homeostasis. Due to its high metabolic activity, it is also the renal compartment more vulnerable to injury. It is well known that the tubular epithelium has regenerative potential; however, this repair capacity is not unlimited and may be dependent on the degree of injury (3). Based on studies from our lab and others, acute injury with proximal tubule death is followed by a wave of tubular proliferation, peaking at 48 hours after injury, to restore tubular cell mass. Lineage analysis indicates that the source of the repairing cells derives from within the tubule rather than a circulating or interstitial progenitor (4). Several lines of evidence indicate that surviving epithelia dedifferentiate, and these dedifferentiated epithelia have an equivalent capacity for repair (5–8). By contrast, a separate body of work has suggested a different model: that a fixed population of Wnt-responsive and/or Pax2-positive intratubular progenitors selectively proliferate and differentiate into proximal tubule cells (9–11).

The phosphatidylserine receptor Kim-1 is induced in acutely injured proximal tubule and binds to apoptotic cells and fragments to clear the tubular lumen of debris (12). Its expression is undetectable in healthy kidney, and expression falls back to baseline after proximal tubule repair is complete (13). Because Kim-1 is not expressed at baseline but is rapidly induced in all injured cells, we reasoned that it cannot mark a fixed progenitor population, and that genetic lineage analysis of injured Kim-1+ cells could address the issue of whether injured, dedifferentiated proximal tubule epithelia are responsible for repair vs. a fixed intratubular progenitor. Importantly, Pax2+ putative intratubular progenitors do not express Kim-1 after injury (10), excluding the possibility that our genetic labeling strategy would include this proposed progenitor population. We created a Kim-1-CreER^t2^ (Kim1-GCE) knockin mouse line, and traced the fate of individual clones labeled soon after injury. We also performed ribosomal pull-down RNA-sequencing during injury and repair to define the repair process in molecular terms. We took advantage of this powerful tool to perform transcriptional profiling to identify the transcriptional signature of the injured tubular epithelial cells. We show that injured, dedifferentiated proximal tubule undergoes proliferative expansion after injury, define transcriptional patterns of these cells during repair, and identify an EGFR-FoxM1 signaling pathway that regulates proximal tubule proliferation after injury.

## Results

### Characterization of mouse model

Kim-1GFPCreER^t2^ (hereafter referred to as Kim1GCE) mice were generated by gene targeting (Supplementary Figure S1). The resulting line knocks out endogenous Kim-1 expression and replaces it with a GFPCreER^t2^ cassette (Figure 1A). To evaluate recombination specificity, bigenic Kim1GCE; R26tdTomato mice received Tamoxifen 6 hours before surgery and on day 1 and 2 after surgery. After unilateral ischemia reperfusion, tdTomato expression was analayzed at days 3 and 14 after surgery (Figure 1B). There was no tdTomato expression at baseline, but in injured kidneys TdTomato expression was localized to the outer segment of the outer medulla. Recombination efficiency at day 3 was unexpectedly low, but there was significantly increased tdTomato expression at day 14 suggesting expansion of the labeled tubular epithelial cells (Figure 1C).

**Figure 1.**
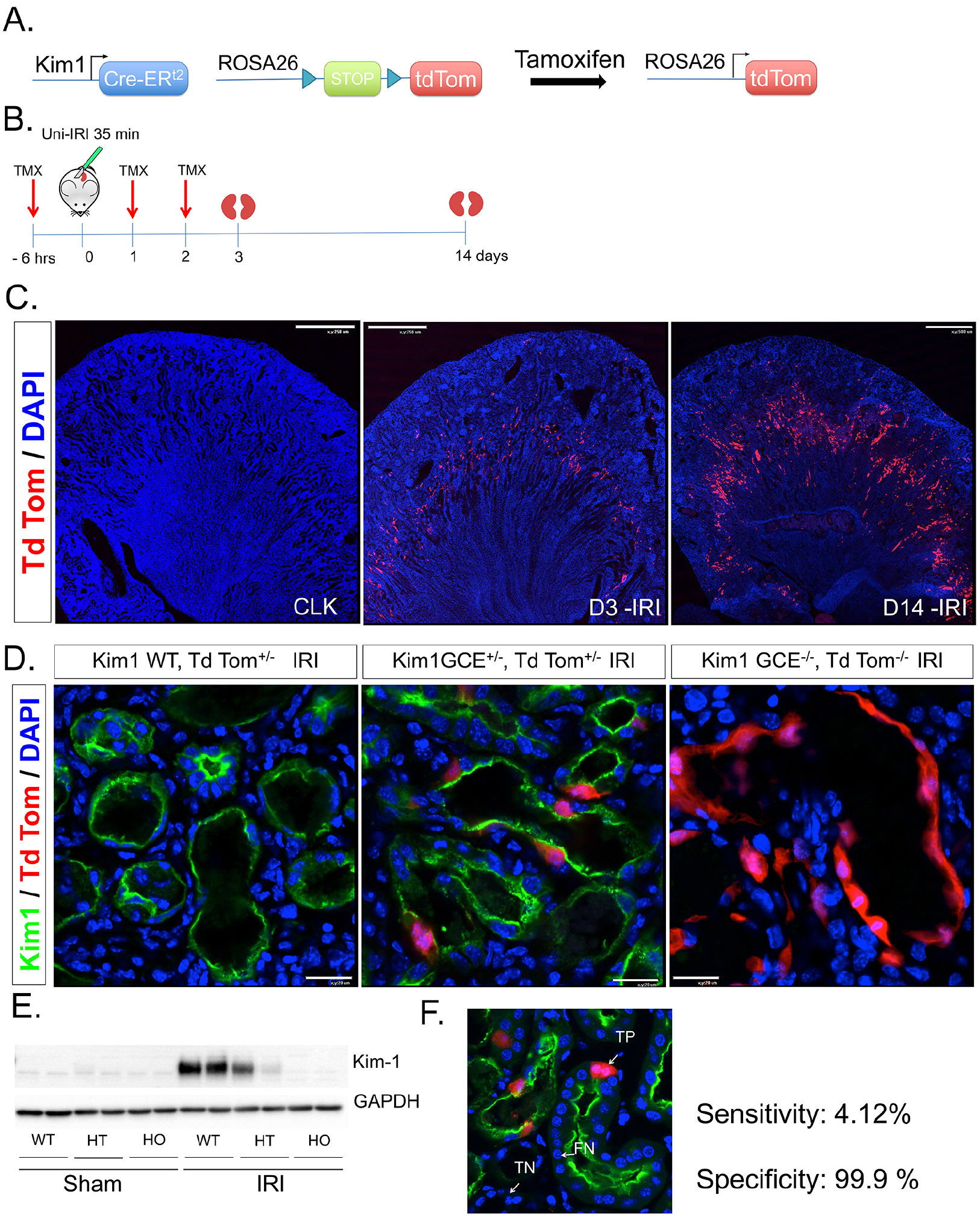
Kim1-GCE Mouse Model. **A.** Kim1-GCE was crossed to the Rosa26tdTomato reporter mouse to allow permanent labeling of injured tubular epithelial cells upon tamoxifen-mediated recombination. **B.** Uni-IRI was performed to validate the mouse model with kidneys harvested at day 3 and day 14 post-injury. **C.** Immunofluorescent staining showing endogenous tdTomato expression in the outer segment of the outer medulla at day 3 with increase expression at day 14. There is absence of tdTomato expression in the contralateral kidney after tamoxifen administration indicating no leaky expression. **D.** Immunostaining with Kim1 antibody showing co-expression with tdTomanto labeled cells in Kim1GCE heterozygous mice. There is absence of Kim1 expression in Kim1GCE homozygous mice as expected since this a knockin to the ATG site. **E.** Western blot for Kim1 showing half the amount of protein expressed in Kim1GCE heterozygous as compared to WT mice and absence of Kim1 protein in Kim1GCE homozygous consistent with the immunofluorescent staining. **F.** Immunostaining showing examples of true positive (TP), true negative (TN), and false negative (FN) for determination of sensitivity and specificity for the mouse model.

To further evaluate recombination specificity, we performed immunofluorescent staining for Kim1. All tdTomato-positive cells also expressed Kim-1 at day 3, although only a minority of Kim-1-positive cells co-expressed tdTomato (Figure 1D). Mice homozygous for the GFPCreERt2 allele did not express Kim1 protein, as expected (Figure 1D and E). To provide a quantitative assessment of the specificity and sensitivity of the model, we counted the number of tdTomato expressing cells that were positive for Kim1 (true positive) and also determined the number of true negatives, false positives, and false negatives as described in the methods section. We determined that the mouse model is 99.9% specific and 4.12% sensitive (Figure 1F). The mechanism behind this low recombination efficiency remains unexplained, however the very high specificity indicates that the line faithfully reports Kim-1 expression without any leaky expression, simply in a minority of cells which appears to be stochastic.

### Lineage analysis reveals clonal expansion of injured proximal tubule after injury

Since Kim-1 is not expressed in healthy kidney, but is induced in all injured proximal tubule very early after injury (rather than a subset), it cannot be a marker of a putative fixed intratubular progenitor cell. We therefore used the Kim-1GCE line to ask whether injured and dedifferentiated cells labeled by Kim-1GCE undergo proliferative repair, or not. Tamoxifen was administered 12 hours after injury at low dose (1 mg) to generate single-cell clones, reducing the possibility of de novo recombination in adjacent cells due to residual tamoxifen (Figure 2A). Kidneys were collected at day 2 and 14 after injury. BUN was measured on day 2, and rose between 80 – 150 mg/dL indicating successful IRI. For the 14-day clonal analysis group, BUN measurement indicated renal recovery as reflected by the reduced BUN from day 2 to day 14 (Figure 2B). Lineage analysis revealed that at day 2 after injury, clones were predominantly single-cell clones in separate tubules but by day 14, there were coherent clones of adjacent tdTomato+ cells (Figure 2C). Careful quantitation revealed that at day 2, 87% of the clones were single-cell. By day 14, the number of single-cell clones had decreased from to 52.2% (p < 0.0005) and the number of multicellular clones (>3 cells) had increased from 2.7% to 31.3% (p < 0.0001, Figure 2D). Maximum clone size was 10 cells, similar to a report tracking Pax2-labeled clones (10). These results indicate that differentiated tubular epithelial cells that become injured are capable of proliferative repair, arguing against the existence of a fixed intratubular progenitor population (6).

**Figure 2.**
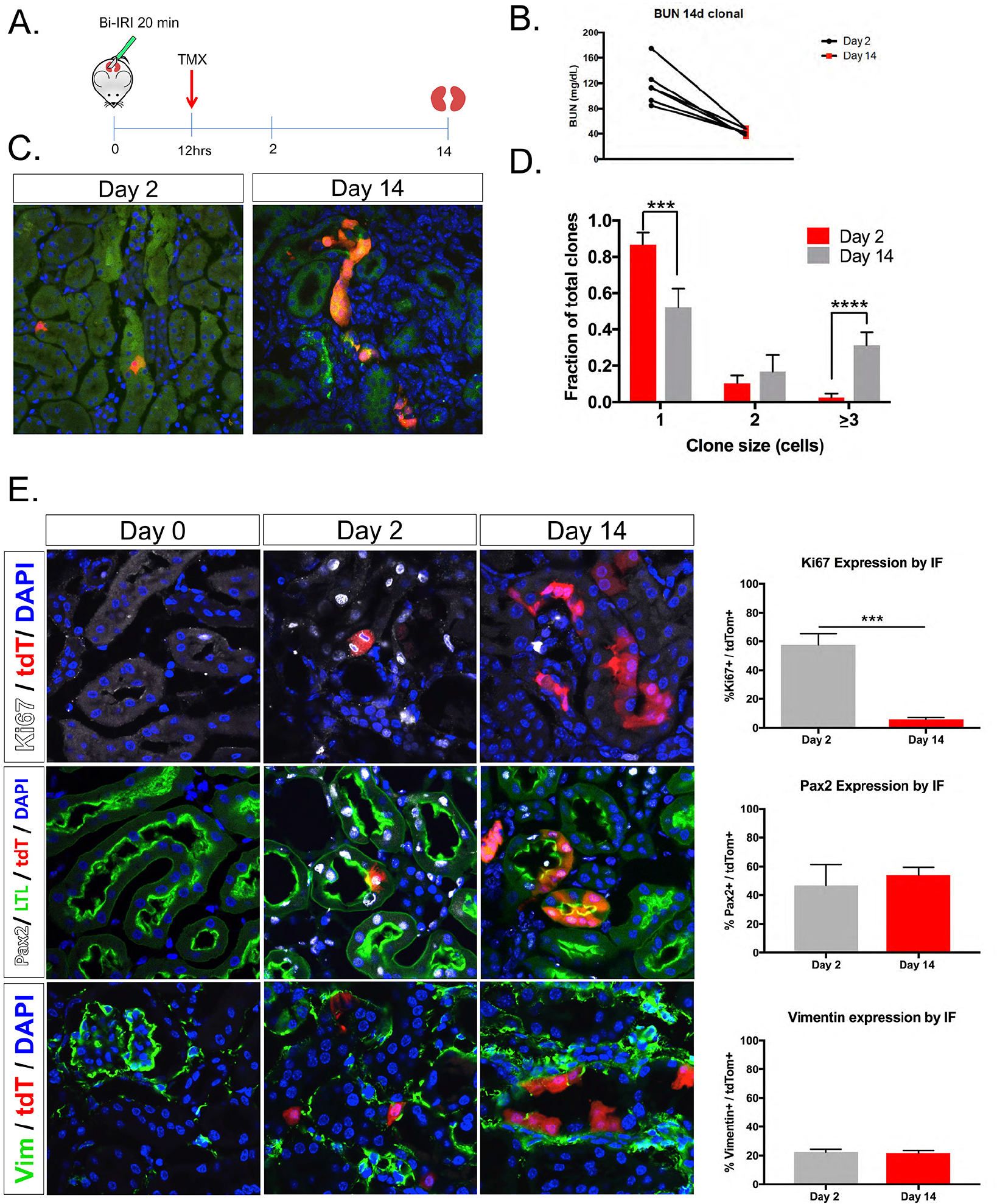
Lineage tracing of injured tubular epithelial cells. A. Kim1GCE;tdTom mice heterozygous for both alleles were subjected to bi-IRI and low dose tamoxifen (1mg) administered 12 hours after surgery. B. BUN increased between 80-150 mg/dL at 48 hours after injury and downtrended at day 14 as expected. C. Immunostaining showing single tdTom cells labeled at day 2 after injury and clusters of tdTom cells at day 14. D. Quantification of clone size at day 2 and day 14 after injury. E. Immunostaining for Pax2, Vimentin, and Ki67 showing coexpression with tdTom cells at day 2. By day 14, there is persistent Pax2 and Vimentin expression in tdTom cells. Ki67 is absent from tdTom cells at day 14 since the cells have completed repair. Quantification showing percentage of co-expression of the tdTom cells with each of the markers.

### Lineage tracing reveals a failed repair population

We next sought to characterize the repair process in more detail. We confirmed that labeled, injured proximal tubule clones undergo a burst of proliferation, because nearly 60% of tdTomato+ cells co-expressed Ki67 at day 2, but only 5% expressed Ki67 at day 14 (Figure 2E). This is in good agreement with reporter of bulk tubular proliferation at this timepoint (4, 14). Pax2 and vimentin are genes that have also been characterized as markers of dedifferentiated proximal tubule cells (14, 15). Two days after injury, we could detect expression of Pax2 and vimentin in about 40% and 20% of tdTomato-labeled cells, respectively. Unlike Ki67, this fraction continued to express these markers at day 14, suggesting some degree of incomplete repair in those populations (Figure 2E).

To investigate further the question of whether tdTomato labeled proximal tubule cells underwent complete repair, or not, we next examined temporal expression of Kim-1 protein and Sox9, which has recently been identified as both a marker of proximal tubule injury and a transcriptional regulator of repair (11, 16). At day 2, about 80% of tdTomato cells co-expressed both Sox9 and Kim1, indicating that these cells are injured and dedifferentiated (Figure 3A and B). This population fell to about 15% by day 14, indicating that while the majority of tdTaomat-labeled cells had successfully repaired, as reflected by their downregulation of Kim-1 and Sox9, about 15% had persistent injury and thus failed repair by day 14. To approach the question from the opposite perspective, we also quantified the number of tdTomato cells that expressed neither Kim1 or Sox9: the population of cells to undergo successful repair. At day 2, these cells were nearly undetectable but by day 14, close to 80% of tdTomato cells were negative for both Kim1 and Sox9 (Figure 3C). These results suggest that injured proximal tubule proliferate after injury, and that the majority have largely completed repair by day 14, but that about 15% remain injured and dedifferentiated at this timepoint, likely reflecting failed repair.

**Figure 3.**
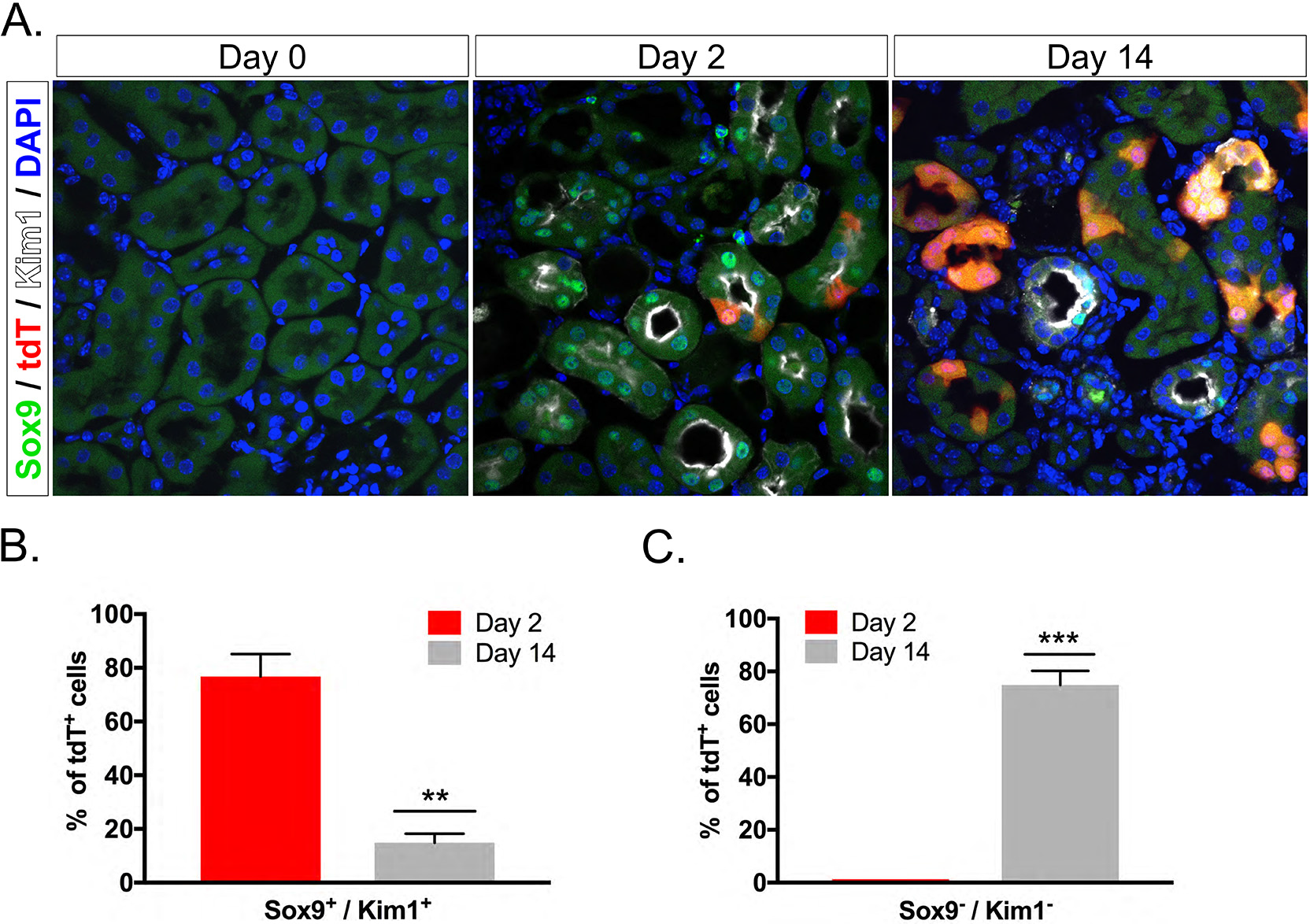
Sox9 immunostaining reveals a small population of cells that have failed repair. **A.** Immunostaining for Sox9 shows absence at baseline (day 0), but expression in tdTom cells upon injury (day 2). At day 14, there are a few tdTom cells that have persistent Sox9 expression suggesting that these are cells that have failed repair. **B.** Quantification of percentage of tdTom cells that express both Sox9 and Kim1 at day 2 and day 14. Quantification of the percentage of tdTom cells that do not express Sox9 and Kim1 at day 2 and day 14.

### Transcriptional Profiling of proximal tubular epithelial cells during injury and repair

Recent work has carefully measured global kidney transcriptional changes over the full course of murine IRI (17). Although this is a powerful resource, it is limited in that relevant cell-specific gene expression signatures may be lost within the integrated expression profiles of the other cell types in the sample. We therefore sought to generate RNA-seq profiles of injured proximal tubule cells during the course of injury and repair by ribosomal pull-down. We generated bigenic heterozygous Kim-1GCE; R26-LSL-eGFPL10a mice in order to perform Translating Ribosome Affinity Purification (TRAP) (18, 19). We isolated mRNA from injured proximal tubule cells at day 2, 7 and 14 after IRI as well as in sham controls. Tamoxifen was administered via gavage 6 hours before surgery and on day 1 after surgery. The isolated polysomal RNA (bound fraction) of 3 biological replicates for each time point was submitted for next generation sequencing. The increased TRAP RNA yield across time points (Supplementary Figure S1B) was consistent with proliferative expansion of labeled cells during repair, which was corroborated by immunofluorescent staining for GFP (Figure 4A). We verified the TRAP protocol by determining the GFP expression by qPCR in the bound vs. unbound fraction since Kim1 is not expressed in uninjured kidney (i.e. sham) (Supplementary Figure S1C). We observed strong enrichment for GFP in the sham and day 2 IRI bound fraction and not in the unbound fraction. The detectable GFP expression in the sham bound fraction is due to leaky eGFPL10a expression in podocytes and collecting duct as previously described (20), but this was about 15-fold lower compared to EGFP expression at day 2 after IRI.

**Figure 4.**
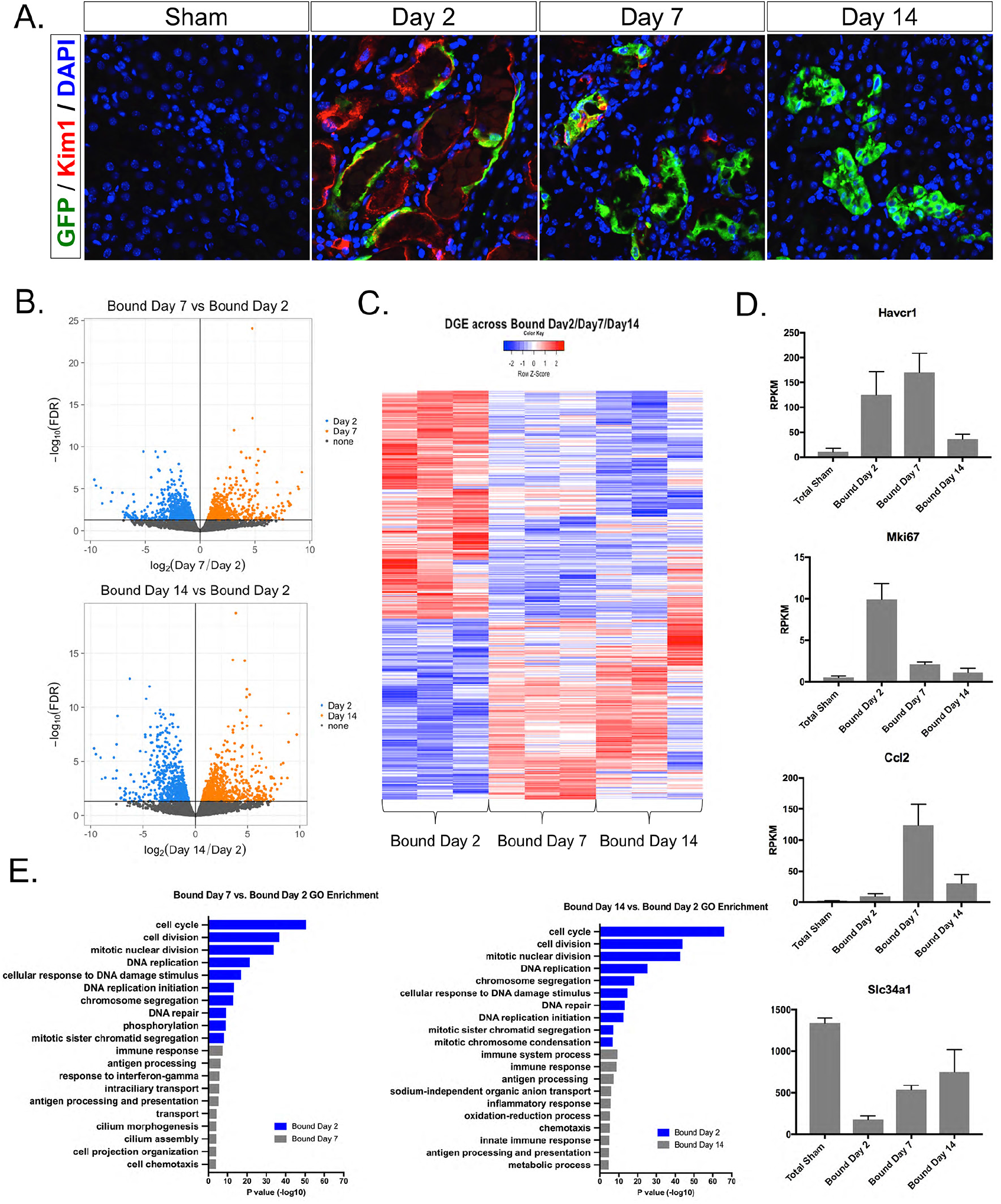
Transcriptional profiling of injured tubular epithelial cells. **A.** Immunostaining for GFP in bigenic Kim1GCE;EGFPL10a kidney sections shows absent GFP expression in sham and co-expression with tdTom cells at day 2 after injury. There is increase GFP expression at day 7 and day 14 since there is clonal expansion of the surviving tubular epithelial cells. **B.** Volcano plots of the DGE list for bound day 7 vs. bound day 2 and bound day 14 vs. day 2. **C.** Heatmap of the DGE list across all 3 time points. **D.** RPKM values across the different time points after injury of known upregulated and downregulated genes. **E.** Gene ontology analysis of the two comparison bound day 7 vs. day 2 and bound day 14 vs. day 2.

We used the edgeR package to perform the differential expression analysis and filtered out the low expressing genes. Principal component analysis (PCA) showed that the biological replicates clustered together across time points indicating a high degree of similarity (Supplementary Figure S2A). The biological replicates for day 14 clustered close to the sham group suggesting that the day 14 group is returning toward the baseline transcriptional state. Since Kim1 is not expressed in sham kidney, we compared the day 7 vs. 2, and day 14 vs. 2 to identify the transcriptional signature during injury and repair. We identified 1457 differentially expressed genes when comparing bound fraction of day 7 vs. day 2 and 1478 differentially expressed genes when comparing day 14 vs. day 2 (Figure 4B). Plotting the DEG list across the 3 time points in a heat map, we observe that about half of the genes are strongly upregulated at day 2 and their expression subsequently falls at day 7 and 14. The other half are genes that are strongly downregulated at day 2, and whose expression rises on day 7 and 14 when the cell is returning to homeostasis (Figure 4C).

To corroborate these results, we selected known markers such as Havcr1 (which encodes Kim1), Ki67, Ccl2 and Slc34a1 during injury and repair and plotted their expression over time points (Figure 4D). These genes reflected the four different patterns we observed. Havcr1 is highly upregulated at day 2 and 7 and decreased by day 14 when the majority of repair is complete, as expected. Ki67 is highly expressed at day 2 after injury since this is the peak of proliferation, but it is much lower at day 7 and 14. Ccl2, an inflammatory marker, is not upregulated until day 7 but then falls by day 14. Finally, the sodium-phosphate exchanger Slc34a1 is significantly downregulated during dedifferentiation at day 2 but expression recovers over time as the tubular epithelial cell redifferentiates.

To dissect further the significance of these differentially expressed genes, we performed DAVID gene ontology (GO) analysis focusing on biological process. The top 10 terms for each comparison (day 7 vs. day 2 and day 14 vs. 2) are shown in Fig 4E. As expected the top GO terms are related to cell cycle and DNA repair since these are key events in the injury response. GO terms for day 7 reflect an immune response and interestingly there were terms related to cilium morphogenesis and cilium assembly suggesting that cilium may play a role during the repair phase. Day 14 GO terms are related to cell transport and metabolic processes suggesting that tubular epithelial cells are returning to homeostasis. We selected some of the most highly expressed genes and performed in situ hybridization (ISH) and qPCR to validate their expression in injured kidney. Candidate genes include: Slc22a7, Rrm2, Ctss, and Sprr2f (Figure 5A and B). Slc22a7 is an organic anion transport with a role in creatinine transport (21, 22). By ISH, we observed that it is expressed at baseline in the S3 segment of the proximal tubule. Upon injury, Slc22a7 is essentially undetectabl, reflecting dedifferentiation. At 14 days after injury, there is re-expression of Slc22a7 since the regenerating cells are now returning to a differentiated state. We observed a similar trend of downregulation during injury and reexpression during repair by qPCR.

**Figure 5.**
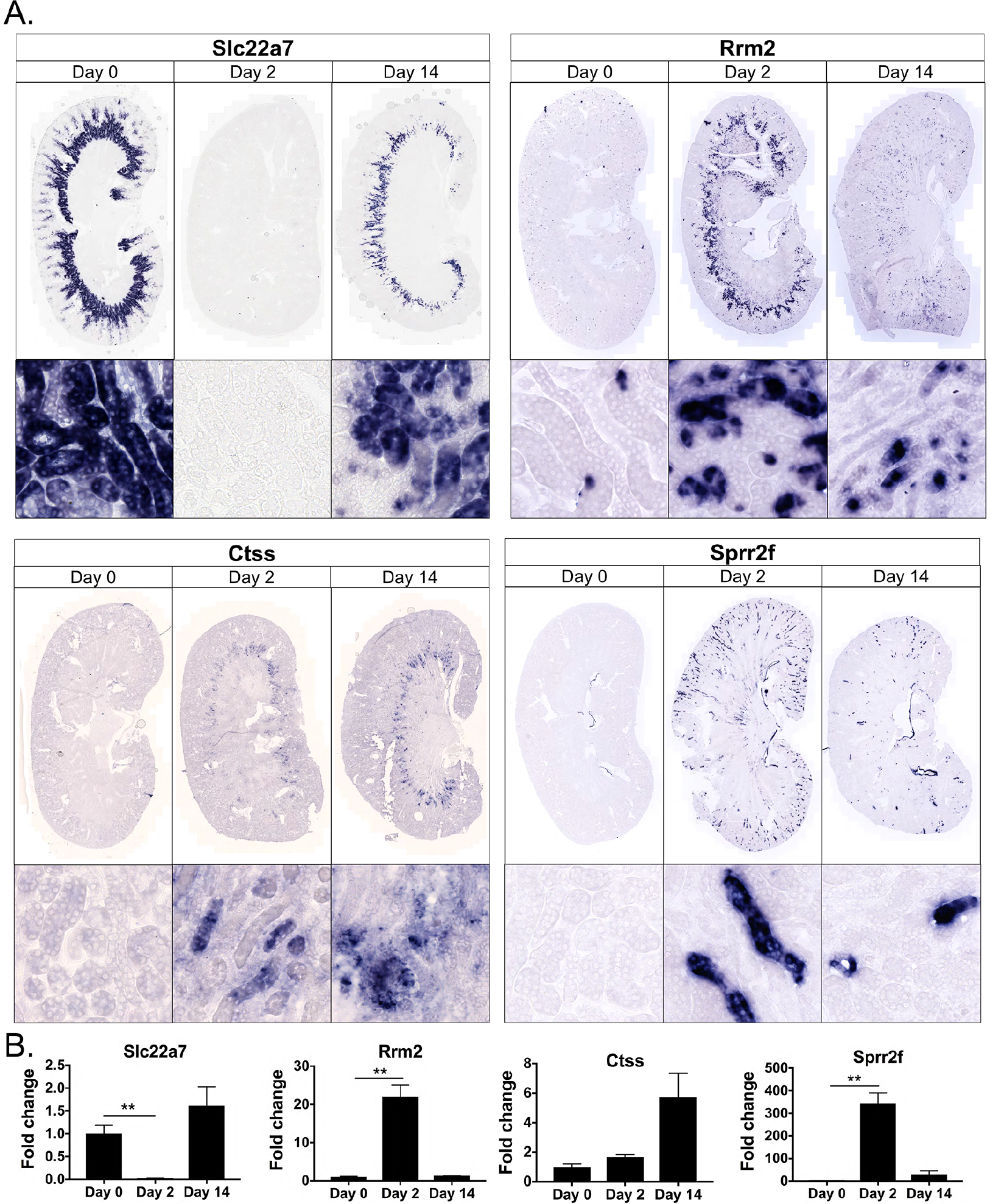
Validation of candidate genes: Slc22a7, Rrm2, Ctss, and Sprr2f. **A.** In situ hybridization in kidney sections from adult, male wild type mouse at 3 different time points after bi-IRI. **B.** qPCR in whole kidney lysates for the candidate genes.

Rrm2 encodes the regulatory subunit of ribonucleotide reductase, which catalyzes the synthesis of deoxyribonucleotides from ribonucleotides (23). By ISH, Rrm2 has a very low level of expression at baseline, and could only be detected in scattered individual tubule cells in the cortex. At day 2 after injury, Rrm2 expression was substantially upregulated, primarily in the outer medulla, which is consistent with the need for DNA synthesis to support cell division in this segment that is damaged the most after IRI. At day 14 post-surgery, Rrm2 expression is decreased and has a pattern similar to that at baseline. Ctss has multiple roles including extracellular matrix degradation and antigen processing and presentation (24, 25). Ctss was not detected in uninjured kidney, but it was expressed in the outer segment of the outer medulla during injury (day 2) and expression even increases by day 14 day as shown by ISH and qPCR. Ctss is involved in EGFR degradation (26); therefore, one can hypothesize that persistent upregulation of Ctss in the tubular epithelium may be preventing further EGFR activation, which could potentially promote renal fibrosis as previously reported (27, 28). Sprr2f belongs to the Sprr family proteins, which are expressed at high levels in the epidermis and function to maintain epithelial integrity (29). At baseline there is complete absence of Sprr2f; however, on day 2 after injury, there is significant expression throughout the cortex, though it is localized to specific tubular segements that are dilated, and thus may represent localized tubular damage. Sprr2f has anti-oxidative functions so could mediate reactive oxygen species detoxification after IRI (30, 31). Fourteen days after surgery, Sprr2f expression is decrease, though individual tubule segments still express it strongly, potentially reflecting failed repair.

### Differential expression of transcription factors and secreted proteins during injury and repair

Transcription factors regulate cell state gene, we therefore identified dynamic transcription factor expression during injury and repair. We took our list of DEG and cross-referenced it against the Riken Transcription Factor Database (32). We compared the lists for Day 7 vs. Day 2 and Day 14 vs. Day 2 and identified 87 and 66 transcription factors respectively. Figure 6A illustrates a portion of the identified transcription factors. Among the transcription factors, we evaluated Ezh2, Foxm1 and Foxj1 in more detail. We also cross-referenced the DEG list for the two comparisons against a database of curated secreted proteins and the results are shown in Figure 6B.

**Figure 6.**
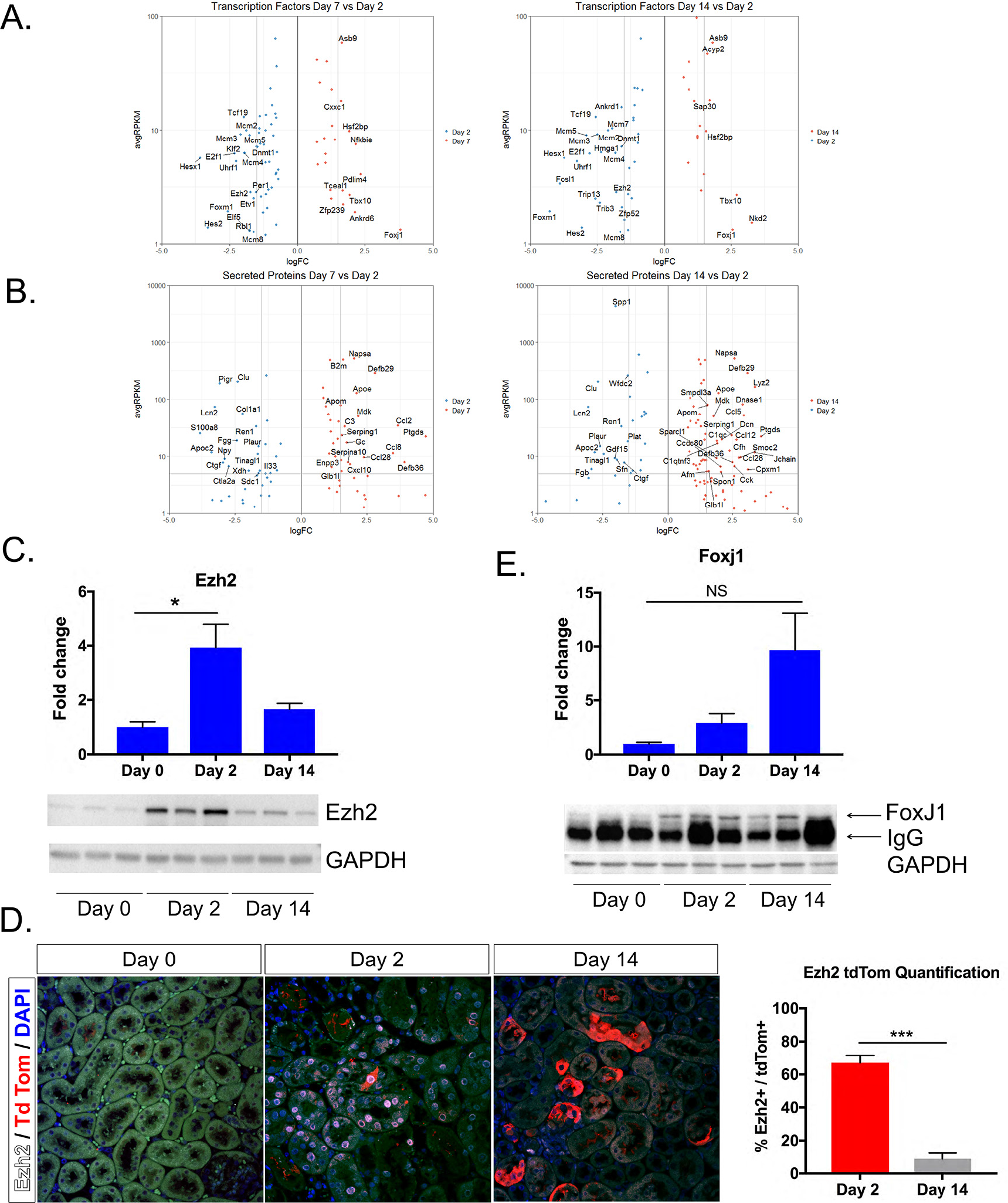
Transcription factors and secreted proteins identified during transcriptional profiling of injured tubular epithelial cells. **A-B.** Scattered plots showing some of the upregulated and downregulated transcription factors and secreted proteins when comparing bound day 7 vs. day 2 and bound day 14 vs. day 2. **C.** Ezh2 mRNA and protein expressions by qPCR and western blot respectively showing upregulation at day 2 and downregulation by day 14. **D.** Immunostaining and quantification for Ezh2 shows co-expression in tdTom labeled cells at day 2 and almost complete absence by day 14 when repair is complete.

Ezh2 belongs to the polycomb repressive complex 2 (PRC2), which participates in methylation of histone 3 (H3K27me) leading to transcriptional repression (33, 34). Ezh2 has been described to have an important role in coordinating cell differentiation in embryonic stem cells (35, 36), mesenchymal stem cells (37–39), hematopoietic stem cells (40, 41), and also in various types of cancer (42, 43). Inhibition of Ezh2 has been reported to prevent renal fibroblast activation (44), though more recently the same group reported that Ezh2 acts in epithelial cells to promote fibrosis (45). We confirmed by qCPR that Ezh2 is upregulated at day 2 after Bi-IRI as compared to baseline (day 0) and its expression downtrends by day 14 (Figure 6C). At the protein level, there is very low expression at baseline (Figure 6C), that was not detectable by immunofluorescence (Figure 6D). However, two days post-Bi-IRI, Ezh2 is highly expressed both by western blot and IF staining (Figure 6C and D). Approximately 65% of tdTomato positive cells also expressed Ezh2. On the other hand, by day 14 after injury, only 10% of tdTomato cells were still expressing Ezh2 (Figure 6D). The temporal pattern of Ezh2 expression suggests that it may be involved with the the transient repression of terminal differentiation genes during injury and repair.

Foxj1 is a transcription factor essential for the assembly of motile cilia (46). Its role in renal injury has been explored in different models in zebrafish, where it was found to be induced upon epithelial injury and required for cilia maintenance (47). We observed Foxj1 upregulation at day 2 after injury with continued expression at day 14 (Figure 6E). By western blot, Foxj1 expression is almost undetectable at day 0 but increased at day 2 and day 14 after Bi-IRI (Figure 6E).

### Foxm1 is upregulated during tubular epithelial injury in murine kidney

One of the most highly upregulated transcription factors in our dataset was Foxm1 which has been considered to be a proliferation-specific transcription factor (48, 49). It is expressed mostly in high cycling organ such as testes and thymus and absent in terminally differentiated cells (48). Foxm1 is also expressed during development in several organs including the kidney and is upregulated in various types of cancers (48, 50, 51). It plays a key role in the G2/M transition and for chromosome segregation and cytokinesis (52) . It also participates in DNA break repair (53). Foxm1 has been found to be reactivated after injury in certain organs such as lung (54), liver (55, 56), and pancreas (57). We sought to validate Foxm1 expression after injury by qPCR and observed that Foxm1 is upregulated 15-fold in day 2 injured kidney compared to day 0 and its expression returns almost back to baseline at day 14 when most of the repair has occurred (Figure 7A). We also evaluated by qPCR downstream targets of Foxm1 related to cell cycle (Ccnb1, Plk1, Aurkb) and DNA repair (Birc5, Brca, Rad51) and found them to be significantly upregulated at day 2 and their expression returning close to baseline at day 14 (Figure 7A). We next designed Foxm1 antisense and sense probes and performed. There was no detectable FoxM1 expression at day 0, but it was clearly expressed in the outer stripe of the outer medulla at day 2, with subsequent downregulation in most (but not all) tubule segments by day 14 (Figure 7B). We next performed ISH on an uninjured and an acutely injured human kidney. FoxM1 was undetectable in healthy kidney, but could be detected in dedifferentiated, flattened epithelia in the acute kidney injury kidney (Figure 7C).

**Figure 7.**
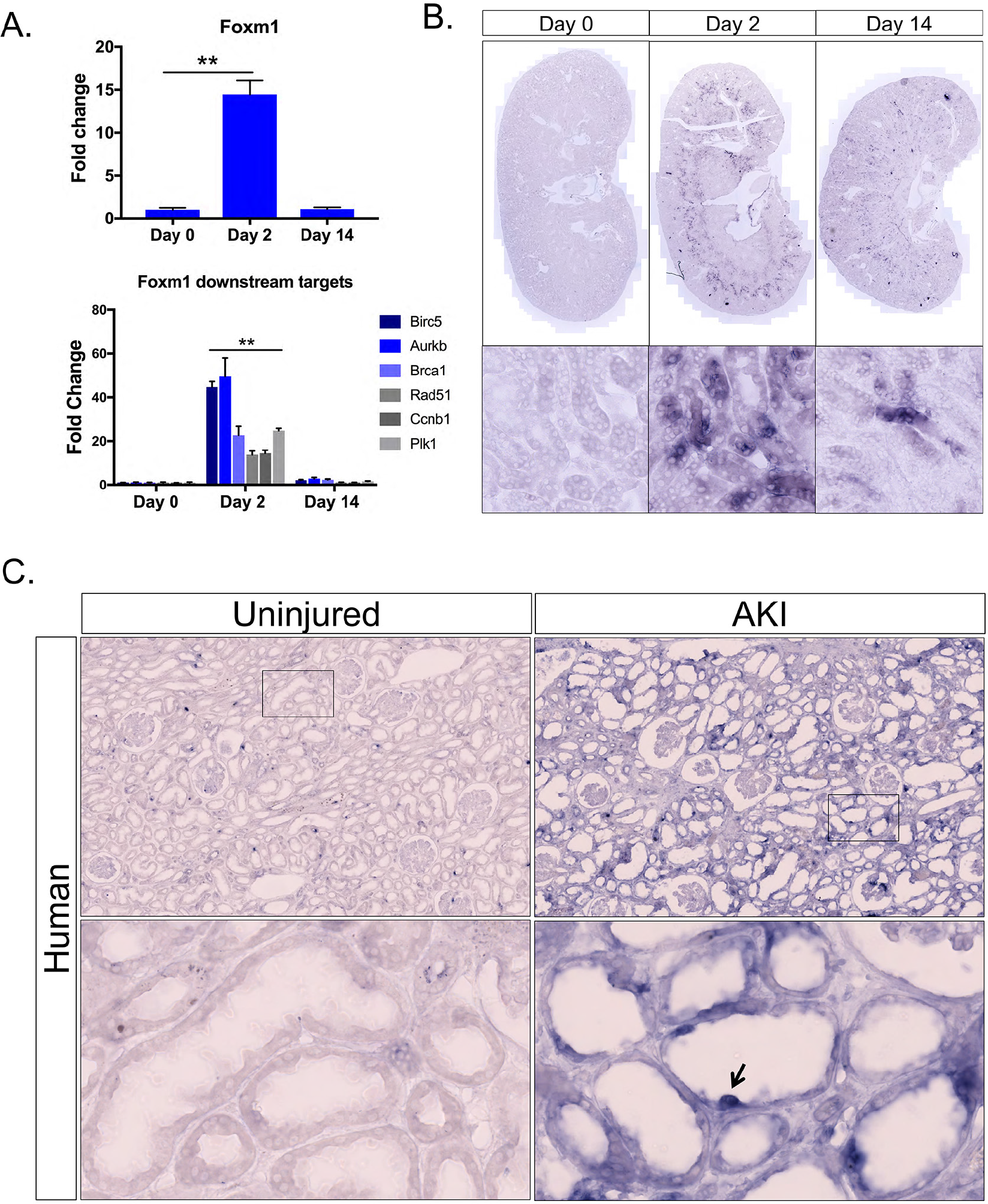
Foxm1 is expressed after kidney injury in mouse and human. A. mRNA expression of Foxm1 and its downstream targets after injury. B. In situ hybridization in uninjured and injured mouse kidneys sections showing increased expression in the outer segment of the outer medulla at day 2 and significant downregulation at day 14. C. In situ hybridization in human samples from uninjured and injured kidney showing absent FOXM1 expression in the uninjured kidney and expression in cells from injured tubules.

### Foxm1 knockdown in human proximal tubular epithelial cells impairs proliferation

Given the known role of Foxm1 in cellular proliferation, we next asked whether the absence of Foxm1 in primary human proximal tubular epithelial cells (hRPTECs) causes a proliferation defect during cell culture. We transfected early passage hRPTECs with Foxm1 siRNA or negative control. Figure 8A shows that Foxm1 siRNA reduced Foxm1 mRNA expression by close to 95% at day 1 and day 2 post-transfection and this was supported by western blot showing absence of the Foxm1 protein in the siRNA treated hRPTECs compared to control (Figure 8B). We evaluated PCNA mRNA expression as a surrogate marker for proliferation and observed that it was downregulated in the Foxm1 siRNA treated hRPTECs, consistent with a proliferative defect (Figure 8C). We also known downstream targets of FoxM1, including cell cycle regulators CCNB1 and PLK1 and DNA repair genes RAD51 and BIRC5. Expression of three out of four of these was reduced with Foxm1 knockdown compared to controls two days after transfection (Figure 8D). We also measured cell proliferation directly. Consistent with the prior results, FoxM1 knockdown hRPTECS had a lower rate of proliferation than the scrambled siRNA controls (Figure 8E).

**Figure 8.**
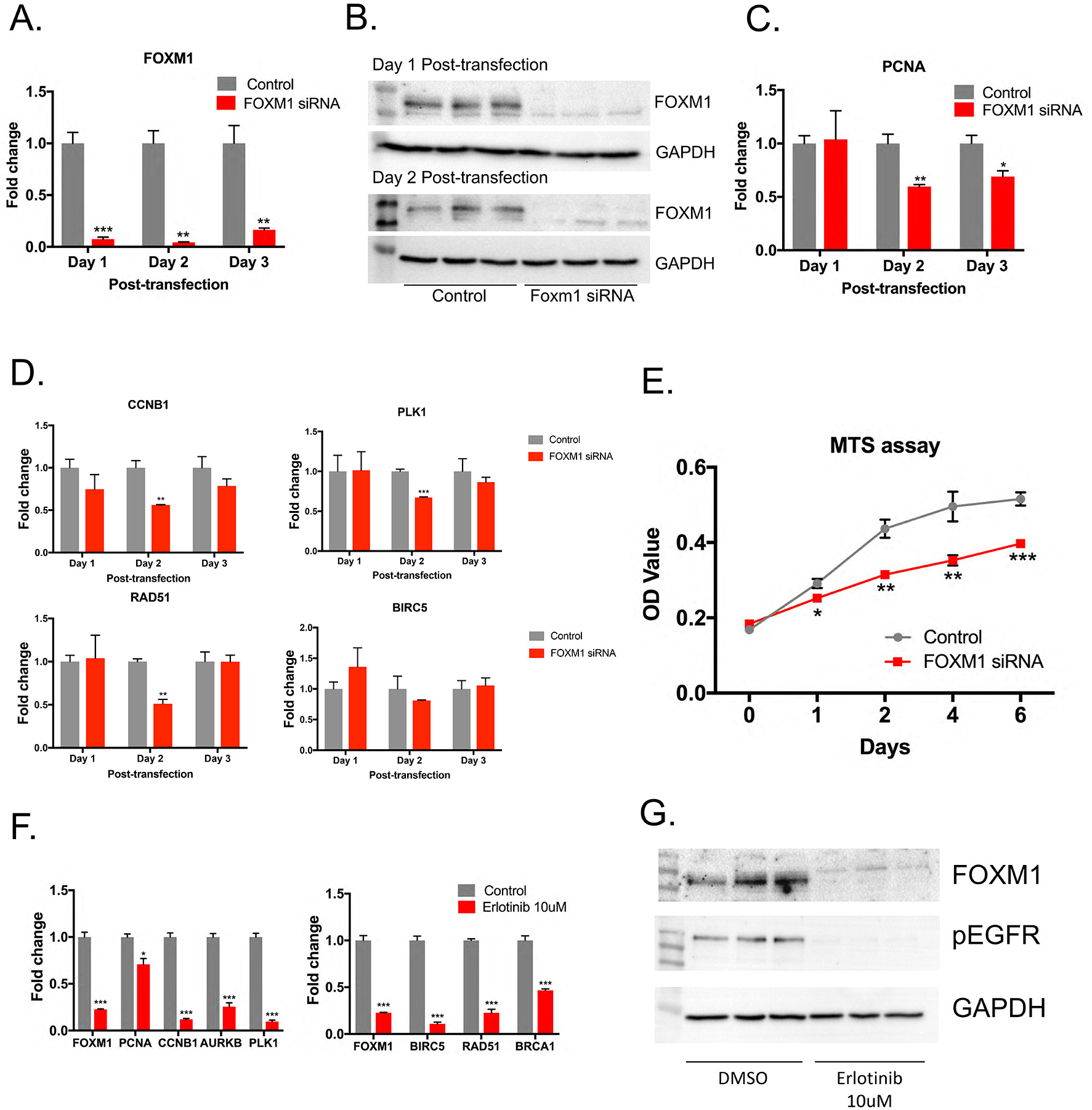
Foxm1 drives proximal tubular epithelial proliferation downstream of the EGFR pathway. A. qPCR for Foxm1 showing efficient Foxm1 siRNA knockdown in hRPTECs at different time points post-transfection. B. Western blot for Foxm1 in hRPTECs corroborating siRNA knockdown. C. PCNA mRNA expression in control and Foxm1 siRNA-treated hRPTECs. D. qPCR for Foxm1 downstream genes in hRPTECs treated with Foxm1 siRNA vs. control. E. MTS assay in hRPTECs shows decrease proliferation in Foxm1 siRNA treated cells compared to control. F. mRNA expression for Foxm1 and several of its downstream targets in hRPTECs after treatment with Erlotinib. G. Western blot in lysates of hRPTECs treated with Erlotinib vs. vehicle. There is complete absence of Foxm1 protein upon inhibition of EGFR with Erlotinib indicating that Foxm1 is downstream of the EGFR pathway. Lack of pEGFR expression confirms inhibition of EGFR by Erlotinib.

### Foxm1 is downstream of the EGFR pathway in human proximal tubular epithelial cells

The EGFR pathway is known to play an important role in tubular epithelial proliferation after injury (58, 59); and FoxM1 regulates keratinocyte cell cycle progression in an EGFR-dependent fashion (60). Therefore, we asked whether Foxm1 expression is regulated by EGFR. We treated hRPTECs with the EGFR inhibitor erlotinib or vehicle. Both Foxm1 mRNA expression as well as FoxM1 downstream targets substantially decreased by erlotinib (Figure 8F). This result was confirmed also by western blot for Foxm1, which showed absence of the Foxm1 protein and absence of the phospho-EGFR protein confirming that the EGFR inhibition was in fact induced (Figure 8G).

## Discussion

There are two primary conclusions from the current study. First, lineage analysis of cells that express Kim-1 after injury shows a proliferative expansion of these dedifferentiated cells during repair. These results do not support a model whereby a fixed tubular progenitor population exclusively repairs injured tubule, because Kim-1 is not expressed in uninjured kidney and it is not expressed in putative Pax2+ intratubular progenitors. Second, we identify an EGFR-FoxM1 pathway that regulates proximal tubule proliferation during repair. Finally, our comprehensive transcriptional analysis of proximal tubule during injury and repair will serve as a resource for understanding dedifferentiation, redifferentiation and failed repair at the molecular level.

It has been proposed that after acute injury, a majority of injured epithelial cells upregulate cell cycle markers, but fail to undergo mitosis – so-called endocycle, and that this explains why past studies relying on cell cycle markers reached incorrect conclusions (10). Our studies here refute this claim, because our genetic labeling strategy excluded putative Pax2+ progenitors since they are resistant to injury and do not upregulate Kim-1 in the first place (10). Rather, we observed clonal expansion of Kim-1-labeled injured cells that indeed completed mitosis, as judged by an expansion in the size of coherent clones during repair.

With respect to our transcriptional analysis of injury and repair, one limitation of our approach is that we do not have a matched baseline proximal tubule-specific transcriptome to compare with, since Kim1 is not expressed in uninjured kidney. As a consequence, we compared acutely injured proximal tubule transcriptomes with those during repair at 7 and 14 days after injury. We recognize that a portion of labeled proximal tubule cells (about 20%) remain injured at day 14. This did not prevent us from defining a large number of transcription factors, ligands and receptors with dynamic expression over the injury and repair timecourse.

During the acute injury phase (day 2) we noted a strong keratinocyte differentiation gene signature. This included the small proline rich (Sprr) genes such as Sprr2f. This family of proteins are induced during keratinocyte differentiation and provide structural integrity to the cornified cell envelope of stratified epithelial (29) . It is unclear what their role might be in tubular repair since tubular epithelium is a simple epithelium. It is possible that Sprr proteins are being expressed as part of the plasticity of tubular epithelium during injury and confer a transient keratinocyte-like phenotype to provide protection in the face of damage and inflammation. Interestingly, some genes from the keratin family were found to be differentially expressed during injury, these included Krt4, Krt5, Krt18, Krt19 and Krt20. Krt18 and Krt20 have been recently recognized as being upregulated after ischemia reperfusion, but their role has not yet been defined (17). Keratins also act as scaffolds that endow epithelial cells with the ability to sustain mechanical and non-mechanical stresses (61); their expression during tubular injury may therefore protect injured epithelia from the harsh postischemic tubular environment.

We identified the transcription factor Foxm1 as strongly upregulated in proximal tubule during injury. Foxm1 is a proliferation-specific transcription factor with expression mostly in high cycling organs such as testes and thymus (48). It is also expressed during development in different organs including the kidney and various types of cancers (48, 50, 51). Multiple studies have provided proof of its role in cellular proliferation and have identified downstream targets that are critical for the G2/M transition and for chromosome segregation and cytokinesis (52). Foxm1 is also re-activated after injury in lung, liver, and pancreas. In a butylated hydroxytoluene (BHT) model of lung injury, Foxm1 was found to be expressed in pulmonary epithelial, endothelial and smooth muscle cells (54). Foxm1 was induced in hepatocytes after liver injury with carbon tetrachloride and partial hepatectomy (55, 56). One study found that absence of Foxm1 lead to impair Beta-cell proliferation after pancreatectomy (57). To date, Foxm1 has only been studied in the context of renal cell carcinoma (62). Based on ISH staining Foxm1 expression localizes predominantly in the S3 segment, which is the area more susceptible to injury, reinforcing the notion that proliferation is an essential aspect in the renal repair response and that Foxm1 may be an important regulator of this process. Our in vitro studies using human proximal tubular epithelial cells confirmed that absence of Foxm1 leads to decreased proliferation, along with downregulation of downstream targets involved in the cell cycle. That EGFR inhibition abolished FoxM1 expression makes sense, since EGF is a potent epithelial mitogen with known importance in regulating the repair response.

In conclusion, surviving, injured epithelial cells are capable of proliferation after injury. Our results do not support the existence of a fixed intratubular progenitor population. We also identify an EGFR-FoxM1 signaling circuit that regulates proximal tubule proliferation after acute injury.

## Methods

### Creation of GFPCreERt2 Knock-in to the *Havcr1* Locus

A targeting vector was constructed to insert the eGFPCreER^T2^-SV40pA (GCE) transgene and a *FRT*-flanked PGKneobpA selection cassette into the 5’ UTR of the *Havcr1* gene. The GCE transgene comprises an enhanced green fluorescent protein (eGFP) and a tamoxifen inducible Cre-recombinase fusion gene (CreER^T2^). A negative selectable marker thymidine kinase (MC1TK) cassette was also included in the targeting vector to select against nonhomologous recombination. The genomic sequence of the mouse *Havcr1* gene and surrounding sequence was downloaded from the University of California, Santa Cruz genome browser (http://genome.ucsc.edu). Repetitive sequences were masked. The lengths of the homology arms were dictated by the repetitive DNA sequence surrounding the target site, resulting in a 1765-bp-kb 5′ homology arm and 2791-bp 3′ homology arm.

A cloning strategy with suitable restriction enzymes and appropriate primers was chosen based on the sequences of the homology arms and all transgene cassettes. Homology arms were amplified from BAC clones containing the *Havcr1* locus, RP23-58M12 and RP23-82L5 (BACPAC Resources Center). using Platinum Pfx DNA polymerase (Thermo Fisher Scientific). All DNA oligonucleotides were ordered from Integrated DNA Technologies. Both homology arms and the transgene cassettes were cloned into pBluescript KS(-). Linearized targeting construct was electroporated into v6.5 (C57BL/6 × 129/Sv F1 hybrid) embryonic stem (ES) cells and transformants were selected by culture in G418 and ganciclovir. Resistant clones were screened by long-range PCR. Positive clones were expanded and underwent injection into albino B6 (C57BL/6J-*Tyr*^*c-2J*^/J) blastocysts using standard procedures.

### Animals

All mouse experiments were performed according to the animal experimental guidelines issued by the Animal Care and Use Committee at Washington University in St. Louis. Kim1-GCE mouse was created as described above. Rosa26tdTomato (JAX Stock # 007909) and C57BL/6J (JAX Stock # 000664) were purchased from Jackson Laboratories (Bar Harbor, ME).

### Surgery

For bilateral IRI, 8-12 week old male mice were anesthetized with isoflurane and buprenorphine SR was administered for pain control. Body temperature was monitored and maintained at 36.5-37.5°C throughout the procedure. Bilateral flank incisions were made and the kidneys exposed. Ischemia was induced by clamping the renal pedicle with a non-traumatic microaneurysm clamp (Roboz. Rockville, MD) for 20 minutes. The clamps were removed at time completion and kidneys returned to the peritoneal cavity. The peritoneal layer was closed with absorbable suture and the flank incisions closed with wound clips.

### Human kidney samples

Human kidney specimens were obtained under a protocol by the Washington University Institutional Review Board. Kidney parenchyma was obtained from discarded human donor kidney with donor anonymity preserved. The injured kidney came from a 62-year-old man with a serum creatinine of 3.3 mg/dL at time of collection. The healthy kidney was from a 38-year-old woman with a serum creatinine of 0.6 mg/dL.

### Translating Ribosome Affinity Purification (TRAP)

Kidneys were harvested and TRAP performed as previously described (19). RNA integrity and quantity were determined using the Agilent RNA PicoChip kit and the Agilent 2100 Bioanalyzer System (Agilent Technologies, Santa Clara, CA). The Clontech SMARTer Universal Low Input RNA kit (Takara Bio USA. Mountain View, CA) was used for cDNA library preparation. cDNA libraries and next generation sequencing were performed at the Genome Technology Access Center at Washington University in St. Louis. Twelve samples were sequenced with an Illumina HiSeq3000, obtaining 25-30 million reads per sample.

### RNA-Seq data analysis

Differential gene expression analysis was performed using the edgeR package (63), and setting a cutoff CPM > 0.4 and FDR of less than 5%. Gene ontology was performed using Database for Annotation, Visualization, and Integrated Discovery (DAVID) (64, 65) and analyzed using the functional annotation tool. Scattered plots for transcription factors and secreted proteins were created using ggplot.

### Real-time PCR

Kidney tissue was snap-frozen in liquid nitrogen at time of harvesting. RNA was extracted using the Direct-zol MiniPrep Plus kit (Zymo. Irvine, CA) following the manufacturer’s instructions. The extracted RNA (600 ng) was reverse transcribed using the High-Capacity cDNA Reverse Transcription kit (Life Technologies. Carlsbad, CA). Quantitative RT-PCR was done using the iTaq Universal SYBR Green Supermix (Bio-Rad. Hercules, CA). Expression levels were normalized to GAPDH and data analyzed using the 2-ΔΔCt method. Primers used are listed on Table X.

### Tissue preparation and histology

Mice were perfused via the left ventricle with ice-cold PBS. Kidneys were harvested and fixed in 4% paraformaldehyde on ice for 1 hour, then incubated in 30% sucrose at 4°C overnight. Next day, tissues were embedded in OCT medium (Sakura Finetek). Kidney sections were cut at 6 μm and mounted on Superfrost slides. Immunofluorescent staining was performed as follows. Kidney sections were washed with 1X PBS for 10 min and permeabilized with 0.25% Triton X for 10 min. Blocking was done with 5%BSA in PBS for 1 hour. Primary antibodies were incubated for 1 hour at room temperature and sections rinsed with 1X PBS for 5 min x 3. Secondary antibodies (1:200) were incubated for 1 hour at room temperature and rinsed with 1X PBS for 5 min x 3. DAPI was used for counterstaining. The following antibodies were used: Kim-1 (AF1817, R&D Systems), Ki67 (14-5698, eBioscience), EGFP (GFP-1020, Aves Labs), Vimentin (Ab92547, Abcam), Sox9 (Ab185230, Abcam), Ezh2 (5246, Cell Signaling).

### Sensitivity and specificity quantification

Kidney sections from Kim1GCE+/−;tdTom+/− were stained with Kim1 antibody (AF1817 R&D Systems), and x400 images (n=10) were taken randomly. A true positive (TP) cell was defined as tdTomato cell that expresses Kim1. A false negative (FN) cell is non-tdTomato cell that does not express Kim1. A true negative (TN) is a non-tdTomato cell that does not express Kim1. A false positive (FP) is a tdTomato cell that does not express Kim1. Sensitivity is then defined as TP/((TP+FN) and specificity as TN/(FP+TN).

### Western blot

Kidney tissue was snap-frozen in liquid nitrogen upon harvesting. Tissue was homogenized in RIPA lysis buffer containing protease inhibitors (Roche). Protein concentration was measured using the BCA assay (Thermofisher). For hRPTECs, cells were washed with 1X PBS, and RIPA buffer added to isolate the cells using a cell scrapper. Using 10% polyacrylamide gel, 10-20 μg of protein was separated by SDS electrophoresis and transferred to an Immobilon PVDF membrane (Millipore). Membrane was blocked with 5% milk in TBST and probed overnight at 4°C with the primary antibody. After washing the membrane with TBST, it was incubated for 1 hour at room temperature with HRP-conjugated secondary antibody (Dako). The membrane was developed using the ECL detection system (GE Healthcare). Primary antibodies: Kim-1 (AF1817 R&D), Ezh2 (5246, Cell signaling, Foxm1 (5436, Cell Signaling), phospho-EGFR (3777, Cell Signaling) and GADPH (A300-641A, Bethyl laboratories).

### In situ hybridization

Kidneys were perfused with RNase-free PBS and fixed with 4% PFA for 1 hour at 4°C and then switched to 30% Sucrose and kept overnight at 4°C. All solutions were prepared in RNase-free PBS. Tissue was embedded in OCT and sectioned into 15 μM sections. In situ hybridization was performed as previously described with some minor modifications (17). Briefly, sections were incubated with 4% PFA at 4°C overnight. After rinsing, sections were incubated with Proteinase K (10 ug/mL) for 20 min, acetylated with 0.375% acetic anhydride and hybridized with sense or antisense digoxigenin-labeled (DIG-labeled) riboprobe (final 500 ng/mL) overnight at 68°C. The next day, sections were washed with a series of stringency washes at 68°C and blocked with 2% Roche blocking reagent for 1 hour at room temperature. Subsequently, sections were incubated with anti-DIG-AP antibody (Roche, 1:4000) at 4°C overnight, followed by development with BM Purple (Roche) which varied from 6 hours to 10 days depending on staining intensity. Finally, sections were fixed with 4% PFA and mounted with Prolong Gold (Invitrogen). ISH images were generated using a Zeiss Axio Scan Z1 slide scanner. DIG-labeled riboprobes were generated from template PCR and the primer sequences are listed in Table X.

### Cell culture experiments

Primary human proximal tubular cells were purchased from ATCC (Manassas, VA) and cultured with Renal epithelium cell growth medium 2 (PromoCell) supplemented with 10 ng/mL EGF, 5% v/v fetal calf serum provided with the medium kit. Cells were maintained in a humidified 5% CO2 atmosphere at 37°C. Experiments were carried out on early passage cells.

### FOXM1 siRNA transfection

hRPTECs were grown to 50-60% confluency at which point they were transfected with 10 nmol/L FOXM1 siRNA (Thermofisher) or negative control siRNA (Thermofisher) using Lipofectamine RNAiMAX (Life Technologies) following the manufacturer’s protocol. Cells were harvested at day 1 and day 2 post-transfection for protein and RNA isolation in order to validate knockdown.

### MTS assay

For the MTS experiments, hRPTECs were transfected with Foxm1 siRNA or negative control as above. One day after transfection, cells were trypsinized (Gibco) and counted using a hematocytometer. Cells were seeded at a density of 1250 cells per well in a 96-well plate in Renal epithelium cell growth medium 2 (PromoCell). Six replicates were prepared per group. Proliferation was measured using the CellTiter 96 AQueous One Solution Cell Proliferation Assay (Promega) as per manufacturer’s protocol. Optical density readings were taken 2 hours after first seeding for day 0, and subsequently on day 1, 2, 4 and 6.

### Erlotinib treatment

hRPTECs were starved overnight by culturing on Renal epithelium cell growth medium 2 without any supplements. Following starvation, hRPTECs were switched to full medium and treated with the epidermal growth factor receptor (EGFR) inhibitor: erlotinib hydrochloride (Cayman chemicals) at concentration 10 uM for 24 hours. Cells treated with DMSO served as control. hRPTECs were then harvested for downstream analysis.

### Statistical Analysis

Data are presented as mean ± SEM. Unpaired t-test was used to compare two groups and p value of less than 0.05 was considered significant. Statistics were performed using GraphPad Prism 7.0 (GraphPad Software Inc, San Diego, CA)

### Mouse Primers

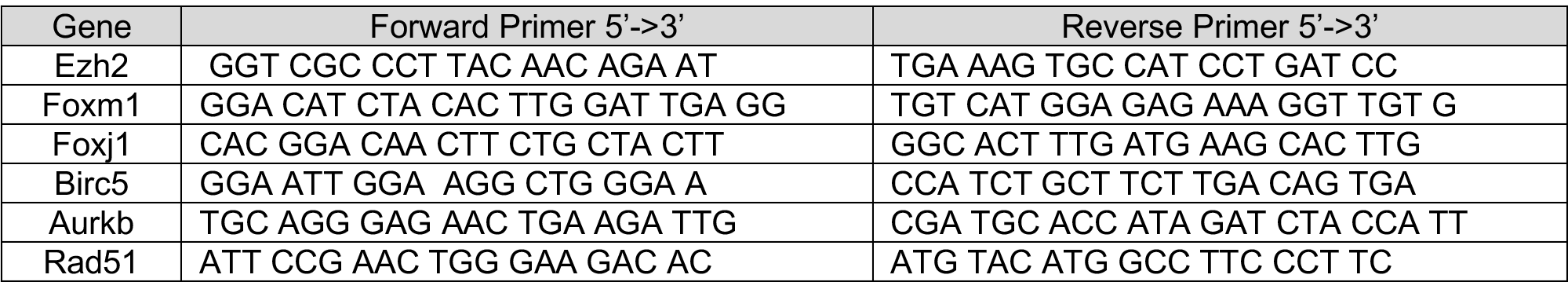

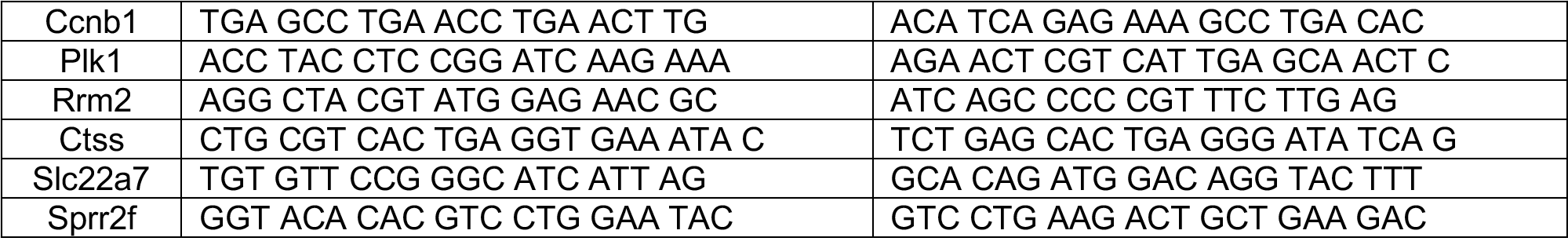

### Human Primers

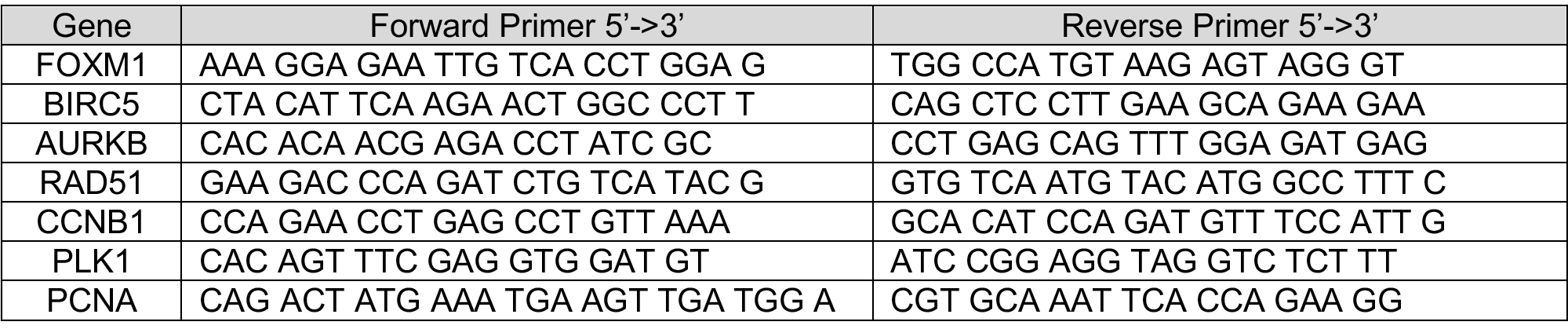

### Mouse Riboprobes

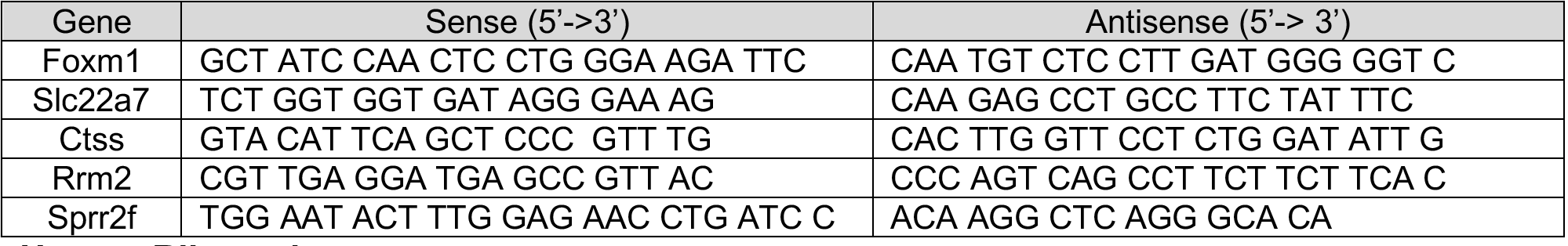

### Human Riboprobe

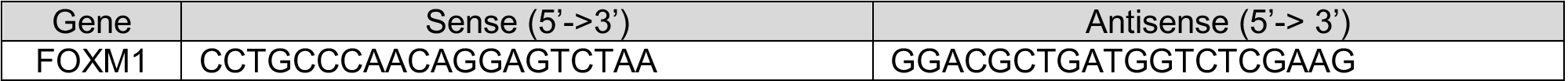

## Author Contributions

MC-P and FFK designed and carried out experiments, analyzed results and reviewed the manuscript. ML, SI, HW and AK carried out some of the experiments, analyzed data and reviewed the manuscript. BDH conceived of the work, designed experiments, analyzed results, and wrote the manuscript with MC-P.

## Acknowledgements

Primary support for this work was from DK107374. Additional support was from NIH/NIDDK grants DK103740 and by an Established Investigator Award of the American Heart Association (all to B.D.H) and by F32 DK103441 (to MC-P).

---

**Supplemental Figure 1.**
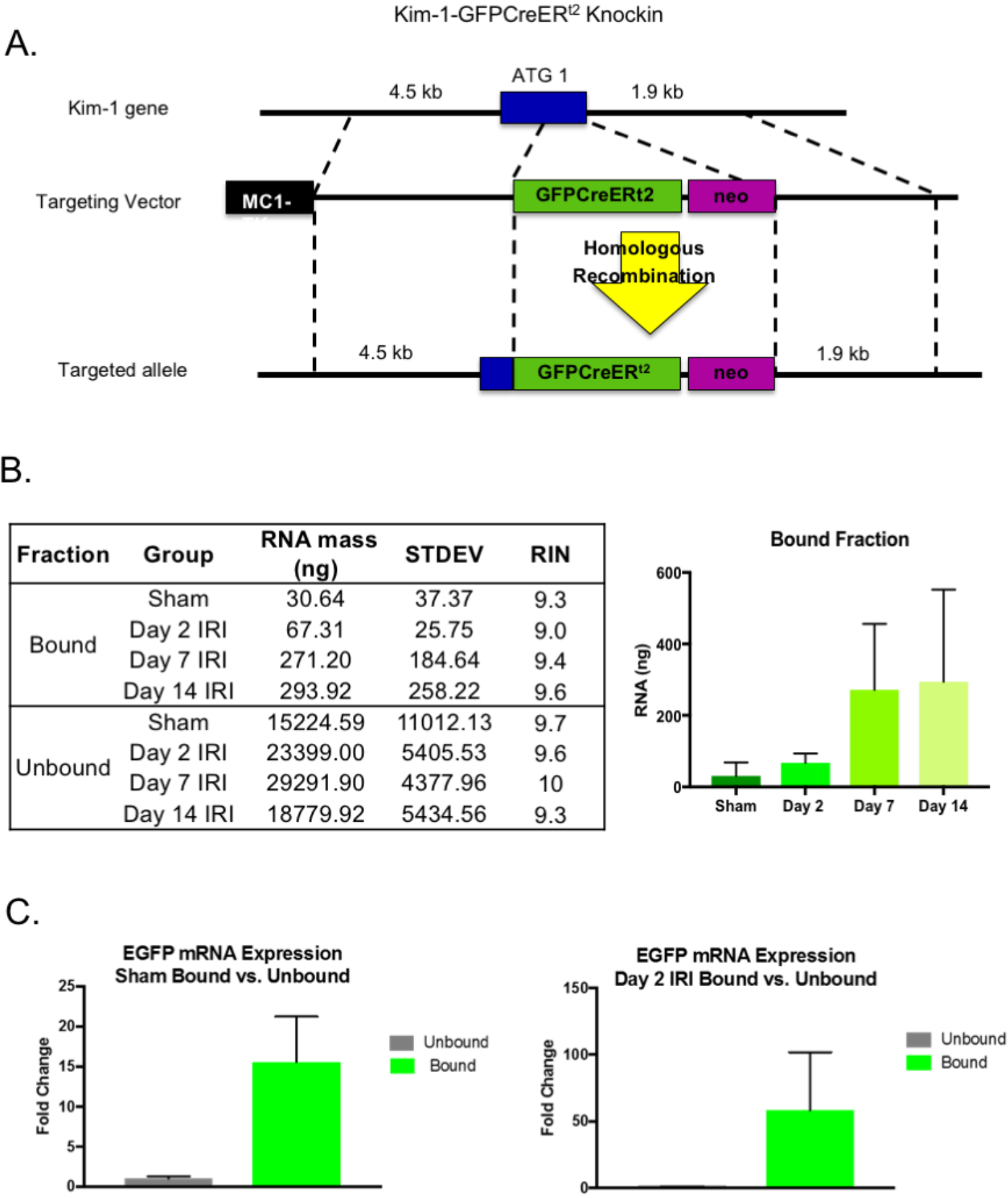
**A.** Targeting strategy design for Kim1-GFPCreERt2 knock-in allele. A targeting vector was used to insert the EGFRPCreERt2 (GCE) transgene and a frt-flanked PGKneobpA selection cassette into the ATG codon of the Havcr1 gene. B. Amount of isolated RNA from TRAP at the different points after injury. C. qPCR for EGFP in bound vs unbound fraction in sham vs. day 2 after injury.

**Supplemental Figure 2.**
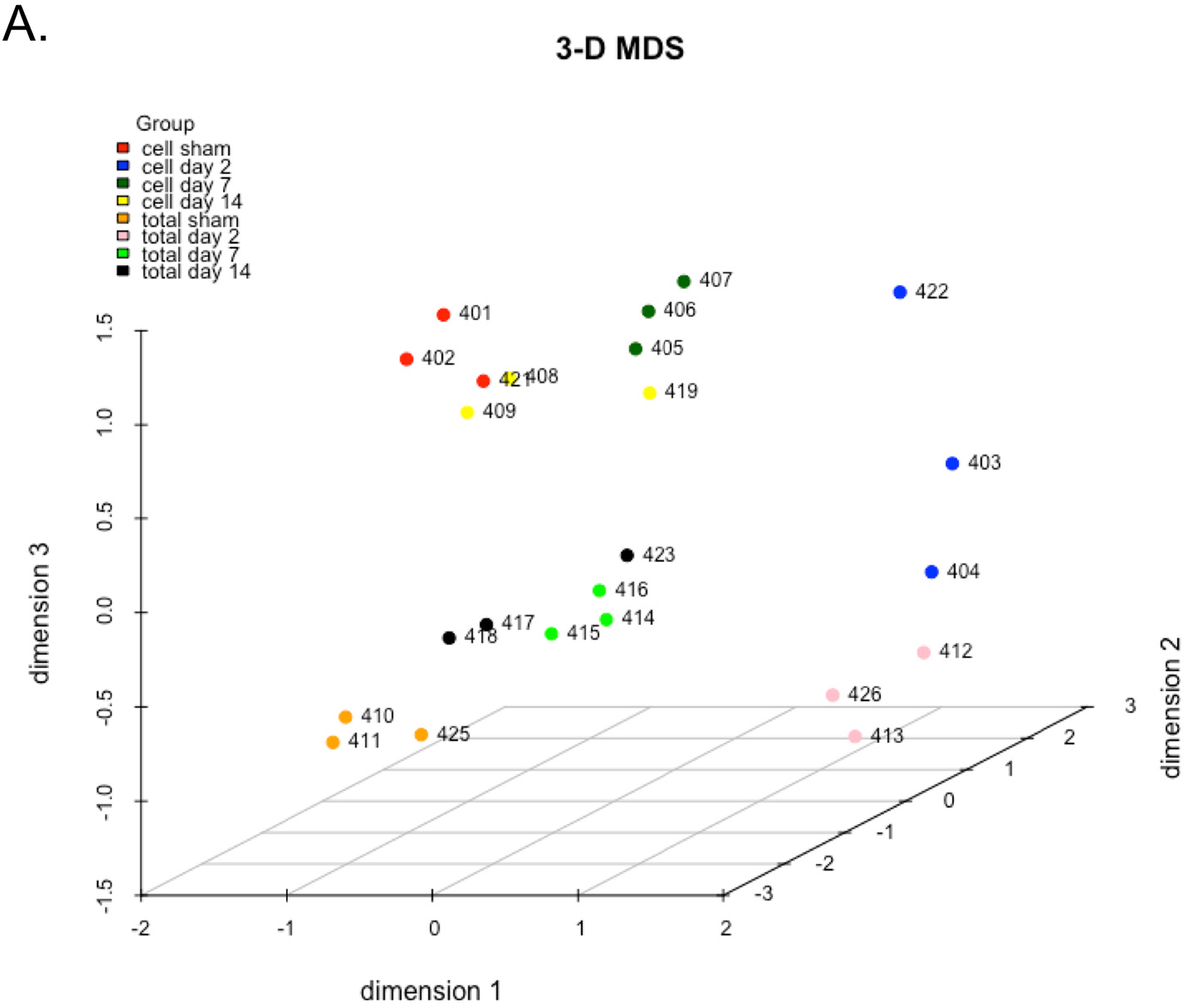
**A.** MDS plot showing good clustering of the samples in cell (bound) vs. total (unbound) at the different time points.

